# Fascicle stretch during partial muscle deactivation does not enhance the human tibialis anterior’s subsequent neuromechanical output

**DOI:** 10.1101/2024.09.25.614882

**Authors:** Brent James Raiteri, Ricardo De Lorenzo, Malte Kraul, Daniel Hahn

**Affiliations:** Human Movement Science, Faculty of Sport Science, Ruhr University Bochum, Bochum, North Rhine-Westphalia, Germany; School of Human Movement and Nutrition Sciences, The University of Queensland, Brisbane, Queensland, Australia

**Author notes:** Correspondence: Brent J. Raiteri.

**Keywords:** dorsiflexor, history dependence, isometric, stretch, torque steadiness

## Abstract

We were interested in whether stretching the muscle via elastic tissue recoil during partial muscle deactivation could trigger stretch-induced mechanisms that subsequently enhance the muscle’s steady-state neuromechanical output. Two torque-controlled experiments were conducted to test this aim. In Experiment 1, fifteen participants performed fixed-end dorsiflexion contractions to a moderate or high-then-moderate level with torque-drop rates of 0, 10, 20, 40, or 80% MVC·s^-1^ from 60-40% MVC, while net ankle joint torque, tibialis anterior (TA) muscle activity level, and TA ultrasound images were recorded. The same measurements were performed in Experiment 2, which tested twelve different participants who performed fixed-end contractions to three reference levels or to a higher-then-lower level (with torque-drop amplitudes of 85-45, 85-30, and 85-15% MVC at 20% MVC·s^-1^). Increased fascicle shortening amplitudes (Experiment 1: 1-2 mm, *p*≤.049; Experiment 2: 5-10 mm, *p*<.001) to initially higher joint torques and different fascicle stretch rates (0.6 to 4.3 mm·s^-1^, *p*≤.041) and amplitudes (4-9 mm; *p*≤.001) did not significantly affect TA’s subsequent muscle activity level relative to the reference contractions at similar joint torques (0-1% MVC, *p*≥.659 and *p*≥.626). However, the torque steadiness relative to the reference conditions was significantly reduced after the 85-15% MVC (*p*=.036) and 85-30% MVC (*p*=.036) torque drops. Consequently, the history of force production affected the control of muscle force more than TA’s steady-state neuromuscular output. These findings indicate that assessing fascicle kinematics concurrently with motor unit behavior during fixed-end contractions with large torque drops might provide unique insights into the neuromechanical contributors to impaired torque control.

**NEW & NOTEWORTHY:** The history of force production affected the subsequent control of dorsiflexion torque more than tibialis anterior’s (TA’s) neuromuscular output following larger amounts of fascicle stretch during fixed-end contractions. Larger fascicle stretches also reduced the median frequency of TA’s EMG signal, which tentatively indicates that the voluntary control of muscle force was altered in the steady state. Together, these findings indicate that voluntary force control might be impaired by stretch-induced and/or activity-level-dependent mechanisms during fixed-end contractions.

## INTRODUCTION

When muscles produce active forces, they shorten or attempt to shorten, which powers or brakes movement, respectively. Despite the importance of active muscle shortening for powering movement, active shortening depresses forces both during and after shortening relative to fixed-end contractions at similar muscle lengths and activation levels (1, 2). Active shortening also increases the energetic cost of force production relative to that in fixed-end conditions and during active muscle stretch at similar lengths and activation levels (3, 4). Conversely, active muscle stretch has the opposite effect. A muscle that attempts to shorten while activated, but that is actively stretched due to an external force that exceeds its force-producing capacity, produces a higher force with a lower metabolic cost than a muscle that achieves similar lengths by actively shortening (2, 5, 6). Intriguingly, this higher force and lower metabolic cost remains even after active muscle stretch ceases during isometric (i.e. steady state) force production (7, 8).

The increased steady-state muscle force following active muscle stretch is typically called residual force enhancement (rFE) (9). However, apparent rFE can also arise after active muscle-tendon unit (MTU) stretch despite concurrent muscle shortening, which we previously speculated to be due to less shortening-induced residual force depression (rFD) rather than stretch-induced rFE and its underlying mechanisms (10, 11) (e.g. increased forces from passive elements (12, 13), altered cross-bridge kinetics (14–16), and/or dissimilarities between half-sarcomere lengths (17)). rFD refers to the long-lasting decrease in force following active muscle shortening (18) and is thought to arise from shortening-induced cross-bridge inhibition (19) or shortening-induced dissimilarities between half-sarcomere lengths (20). We postulated that rFD was reduced after MTU stretch because of less muscle shortening during active force development in MTU stretch-hold versus fixed-end reference conditions. Less shortening occurred during MTU stretch because this maneuver effectively reduces series compliance (21). Consequently, less shortening-induced rFD might lead to ‘apparent’ rFE (10, 11, 21–23).

As fixed-end contractions can be contaminated by rFD (10), and rFD is thought to be triggered by unimpeded muscle fiber shortening under activation (10, 19, 20), we wanted to explore whether rFD can be reduced by muscle (i.e. fascicle) stretch during fixed-end contractions. This is because active muscle stretch and thus sarcomere stretch is thought to trigger rFE-based mechanisms (9, 24, 25). rFE was typically observed following simultaneous muscle stretch and MTU stretch in EMG-controlled experiments *in vivo* (26–28) and during artificially-induced contractions (29–34). When torque instead of muscle activity level is controlled, rFE was observed indirectly as a reduction in the agonist muscle’s activity level following MTU stretch relative to reference conditions without MTU stretch (35–37). However, whether rFE is present after fascicle stretch from partial muscle deactivation without simultaneous MTU stretch remains to be systematically evaluated among different fascicle stretch conditions.

We previously tested whether a force drop at one fixed rate and amplitude could reduce the muscle activity level at a given force during fixed-end dorsiflexion contractions (11). However, we found no significant difference in the tibialis anterior (TA) muscle activity level after fascicle stretch compared with a reference condition without fascicle stretch in torque-controlled experiments (11). We also found no significant difference in the steady-state force at a given muscle activity level following fascicle stretch compared with a reference condition without fascicle stretch in TA EMG-controlled experiments (11). Earlier findings from the adductor pollicis somewhat agree with our findings, except that some participants exhibited enhanced steady-state forces of 5% and 21% following respective force drops from 60-30% and 100-30% during EMG-controlled experiments (33). However, more recent findings are contradictory, which included a higher muscle activity level and greater torque variability following versus preceding a triangular ramp contraction that was superimposed in the middle of fixed-end dorsiflexion contractions during torque-controlled experiments (38). These more recent findings rather indicate that the higher muscle activation preceding the force drop, which might induce greater rFD, reduced the neuromuscular system’s capacity to produce and control a subsequently lower steady-state force under voluntary drive.

From the available data, it is unclear whether fascicle stretch due to partial muscle deactivation enhances, depresses, or does not change a muscle’s subsequent steady-state neuromechanical output during fixed-end contractions. Therefore, we aimed to test whether different fascicle stretch rates (Experiment 1) or fascicle stretch amplitudes (Experiment 2) affected TA’s subsequent steady-state EMG amplitude (i.e. muscle activity level) following partial muscle deactivation in torque-controlled experiments. Torque was controlled rather than activity level because torque variability during a contraction is lower and thus easier to match to a predefined trace (11). We also included reference conditions within the experimental protocol that involved ramp-hold contractions with constant steady-state torque production (i.e. no torque drop during the contraction) to investigate whether increasing fatigue contributes to a potential increase in TA’s muscle activity level. We expected that a faster drop in torque and correspondingly faster rate of fascicle stretch would not progressively change TA’s subsequent steady-state muscle activity level, but that this muscle activity level would be reduced compared with a reference condition without fascicle stretch. We also expected that a larger drop in torque and correspondingly larger fascicle stretch amplitude would lead to greater relative reductions in TA’s steady-state muscle activity level compared with reference conditions. These expectations were formulated based on two major assumptions: 1) that additional rFD relative to the reference condition would be induced by increased muscle activity coupled with additional fascicle shortening (19, 39, 40), and; 2) that fascicle stretch during partial muscle deactivation would be active for the fibers that were not de-recruited, which would trigger rFE-based mechanisms within them to subsequently more than offset the additional rFD from 1). As rFE is independent of stretch speed at slow and moderate velocities (2, 41, 42), but dependent on stretch amplitude at longer muscle lengths (39, 43), we had different *a-priori* expectations for the experiments.

## MATERIALS AND METHODS

### Participants

Fifteen (Experiment 1: age: 26±3 yr (mean±standard deviation); mass: 72±14 kg; height: 176±11 cm) and twelve (Experiment 2: age: 25±2 yr, mass: 74±9 kg, height: 178±7 cm) healthy and recreationally-active sport science students (7 and 3 women, respectively) were recruited from the Faculty of Sport Science at Ruhr University Bochum, who provided free written informed consent prior to participating in either study. Both study protocols were conducted in accordance with the Declaration of Helsinki and were approved by the Faculty of Sport Science’s local ethics committee. G*Power (v3.9.1.7 (44)) was used to determine that a minimum sample size of ten was required to achieve at least 80% power to detect a minimum effect size of interest (*d*z) of 1 between two conditions of interest with a two-tailed alpha level of 5%. Thus, with these sample sizes, only large effects between two paired conditions could be detected.

### Experimental setup

Participants were secured prone on a bench via a lashing strap while they performed voluntary contractions with their right dorsiflexors against a motorized dynamometer. The participant’s right foot was secured to the dynamometer’s footplate attachment by a custom-built adjustable U-shaped frame that was fixed over the metatarsals, as well as a Velcro strap over the mid-foot. This setup limited joint rotation during dorsiflexion contractions, as well as force contributions from the toe extensors to the measured net ankle joint torque. The heel of the participant’s right foot was also flush against the footplate’s rigid heel support, which prevented the foot from translating on the footplate, and foam was positioned between the heel and footplate to minimize discomfort. The participant’s right ankle joint was fixed for the duration of the experiment at an angle of ∼10° plantar flexion, where 0° represents a perpendicular angle between the shank and sole of the foot. The tested angle was verified during a 50% perceived effort contraction with the help of a digital goniometer (EPT-DAF 380 mm, Conrad Electronic SE, Hirschau, Germany). Ankle joint misalignment was minimized during the 50% perceived effort contraction by manipulating the dynamometer’s axis of rotation to ensure that a laser pointer, which was projected from this axis, was located over the lateral malleolus of the right foot.

The experimental techniques used are similar to those already described in Raiteri et al. (45) and will only be briefly described below. The data collection system (Cambridge Electronic Design Ltd, Cambridge, UK) used a ±5 V input range and a 16-bit analog-to-digital converter (Power1401-3) coupled with Spike2 software (64-bit version). This system temporally synchronized all recorded digital signals. Net ankle joint torque and crank arm angle were measured at 2 kHz via dynamometry (IsoMed2000, D&R Ferstl GmbH, Hemau, Germany). TA’s muscle activity level was recorded as a single differential signal at 2 kHz via electromyography. Surface electrodes (hydrogel Ag/AgCl, 8 mm recording diameter, H124SG, Kendall International Inc, Mansfield, Massachusetts, United States) were attached to the skin above TA’s distal superficial compartment in a bipolar configuration (2 cm inter-electrode distance), as well as above the left lateral femoral condyle (ground electrode in Experiment 1) or right fibular head (ground electrode in Experiment 2), following standard skin preparation. TA muscle architecture changes were visualized via PC-based brightness-mode ultrasound imaging (LS128 CEXT-1Z, TELEMED, Vilnius, Lithuania) and a coupled flat, linear-array transducer (LV7.5/60/128Z-2). The transducer was secured over TA’s mid-belly just proximal to the bipolar electrodes with self-adhesive bandage. Ultrasound images were captured at ∼34 fps with a frequency of 6 or 8 MHz and a 60 mm (width) × 50 mm (depth) field of view.

### Experimental protocol

Participants in either experiment performed the same protocol over two sessions. The familiarization session occurred on a separate day prior to the experimental session. The dorsiflexor MTUs were first preconditioned with five submaximal voluntary (80% perceived effort, 1-s hold, 1-s rest) dorsiflexion contractions and one near-maximal (∼95% perceived effort, 3-s hold) contraction (46). After two minutes of rest, participants then performed two to four maximal voluntary contractions (MVCs; 3-s to 5-s hold) until the peak-to-peak torque was no longer increasing in the subsequent contraction and within 5% between two contractions. Participants were told by the investigator to pull the top of their right foot towards their shank as hard as possible using only their dorsiflexor muscles and they received real-time visual feedback of their net ankle joint torque via a screen positioned in front of them. At least two minutes of rest was provided to minimize fatigue following each MVC.

Once the highest MVC torque was determined, participants attempted to match their real-time active torque (i.e. passive net joint torque at rest subtracted from the recorded net joint torque) during fixed-end dorsiflexion contractions to within predefined traces that were 6% MVC (Experiment 1) or 8% MVC (Experiment 2) apart. The mean desired torque trace for each condition in both experiments are shown in Figure 1. In Experiment 1, the ascending ramp level was either moderate (40% MVC) in a reference (“Ref”) condition, or high (60% MVC; Hold 1) in four so-called “High-Mod” conditions. After the high level was achieved in the High-Mod conditions, the active torque was then reduced to the same moderate level as in the Ref condition (40% MVC) and then maintained (Hold 2). The descending ramp rate from 60-40% MVC varied between High-Mod conditions and was “Slow” (2.00 s), medium (“Med”; 1.00 s), “Fast” (0.50 s), or “Rapid” (0.25 s). In Experiment 2, the ascending and descending ramp rates were kept constant among conditions, but the ramp levels varied. The ascending ramp level was either minimal (“Min”; 15% MVC), “Low” (30% MVC), or moderate (“Mod”; 45% MVC) in three reference conditions, or close to maximal (85%; Hold 1) in three so-called “Max” conditions. A higher normalized Max torque (e.g. 90 or 95%) was not used because pilot testing revealed that this level was too high to consistently match for the duration of the experiment. After the Max level was achieved in the Max conditions, the active torque was then reduced to the same Min (“Max-Min”), Low (“Max-Low”), or Mod (“Max-Mod”) level as in each reference condition and then maintained (Hold 2).

**Figure 1.**
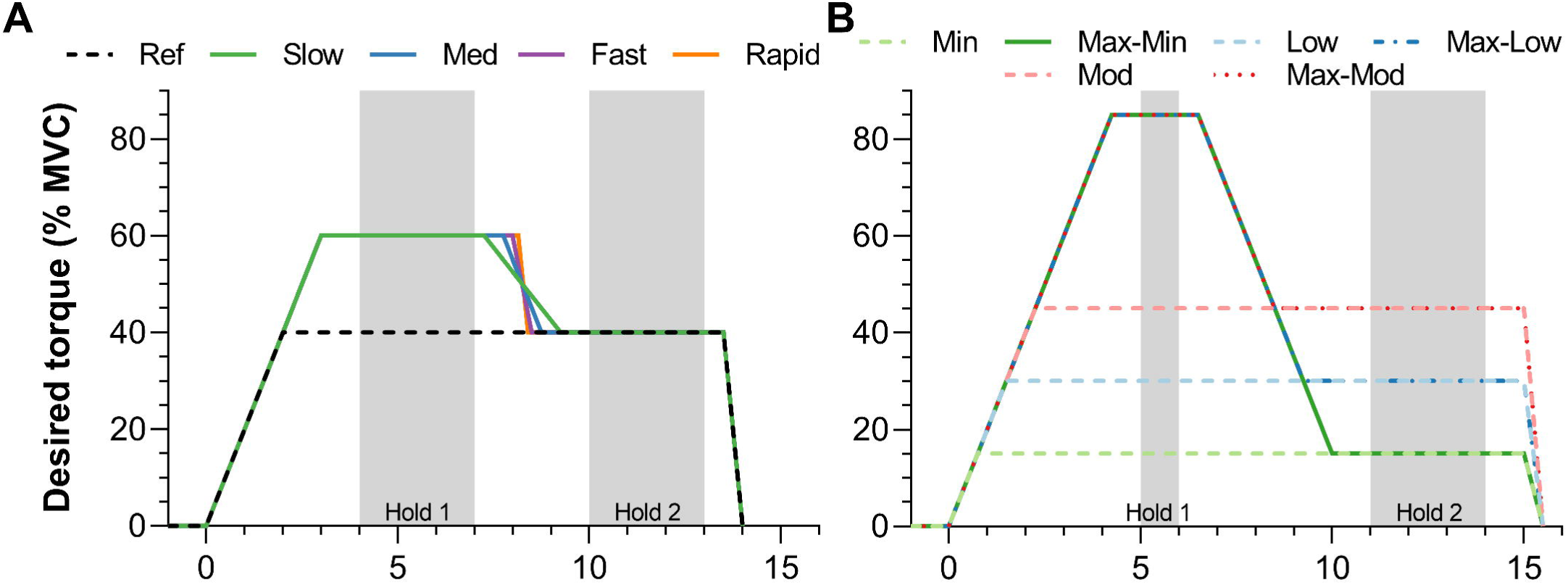
Desired active torque traces that participants attempted to match during fixed-end dorsiflexion contractions in (A) Experiment 1 and (B) Experiment 2. All contractions lasted 13.5 s in Experiment 1 and 15 s in Experiment 2. The ascending and descending ramp rates were fixed at 20% of maximum voluntary torque (MVC)·s^-1^ and variable in Experiment 1, respectively, whereas both rates were fixed at 20% MVC·s^-1^ in Experiment 2. The absolute difference in active torque between the measured and desired trace from contraction onset to the end of Hold 2 needed to be less than 10% of the maximum voluntary torque at 10° plantar flexion for the trial to be considered valid. The grey shaded areas indicate the two analyzed steady-state phases (Hold 1 and Hold 2) of the contractions.

In Experiment 1, conditions were performed in blocks (i.e. five trials per block because there were five conditions), the order of conditions in each block was randomized, and at least four blocks (i.e. four contractions per condition) were performed. In Experiment 2, the Mod condition was executed first and last in the first and third blocks to check for fatigue, as well as in a randomized location in the second block. The order of the other conditions in each block was randomized and at least three blocks (i.e. three contractions per condition) were performed. Within each block, a High-Mod (Experiment 1) or Max (Experiment 2) trial was deemed invalid if the torque deviated 10% MVC or more from the desired torque trace, and invalid trials were immediately repeated up to three times. If the fourth consecutive trial was also deemed as invalid, then that condition was postponed and repeated after the last block was completed. At least one minute or two minutes of rest separated the trials and blocks in Experiment 1, respectively, and due to the higher contraction intensities involved in Experiment 2, at least two or three minutes of rest separated the trials and blocks, respectively.

### Surface electromyography

Electromyography was performed using different systems in the two experiments (Experiment 1: AnEMG12, OT Bioelettronica, Torino, Italy; Experiment 2: NL 844 Pre-Amplifier and NeuroLog System, Digitimer Ltd, Hertfordshire, UK). Regardless of the system, EMG signals were amplified 1000 times and band-pass analog filtered between 10-500 Hz. Prior to electrode placement, ultrasound imaging was used to verify that the bipolar electrodes over the TA were in line with its muscle fascicles, as well as away from the muscle’s borders at rest and during contraction.

### Ultrasound imaging

A flat linear-array transducer was centered over the TA mid-belly in the sagittal plane in a custom-made polystyrene frame to minimize local muscle compression and discomfort. The transducer’s imaging face was coated in water-soluble transmission gel and secured over the skin once the fascicles and aponeuroses of TA’s superficial and deep compartments were clearly visible.

### Data analysis

Data were processed offline using custom-written scripts in MATLAB (R2022a 64-bit version, MathWorks, Natick, Massachusetts, United States). First, the recorded digital signals from each trial were cropped between a common start and end time using the ultrasound timestamps and then exported as .mat files and combined with tracked fascicle data as previously described (10). The tracked fascicle data included absolute lengths of a representative fascicle within TA’s superficial compartment, which was tracked automatically by applying the Lukas-Kanade-Tomasi algorithm using an updated version (https://github.com/brentrat/UltraTrack_v5_3) of UltraTrack (47).

Dynamometer data were filtered using dual-pass second-order low-pass Butterworth filters with corrected cut-off frequencies (48) of 20 Hz for net joint torque data and 6 Hz for crank arm angle data. Active torque was calculated by subtracting the filtered passive torque from the filtered recorded torque of each trial. Passive torque was calculated as the mean filtered recorded torque before contraction onset over 1 s (all submaximal voluntary contractions), 0.5 s (MVCs from Experiment 1), or 0.25 s (MVCs from Experiment 2). TA’s muscle activity level was calculated as follows: first, the raw EMG signal was bandpass filtered between 20-400 Hz with a dual-pass second-order Butterworth filter; second, the mean bias was subtracted; third, this corresponding signal was rectified, and; lastly, the rectified signal was smoothed with a dual-pass second-order low-pass Butterworth filter with a corrected cut-off frequency (48) of 10 Hz. Active torque and muscle activity level data (0.5-s window centered on the maximum amplitude) were then normalized to the corresponding values attained during the MVC trial with the highest active torque.

Fascicle shortening amplitude was calculated as the maximum difference between fascicle lengths from 0.5 s before contraction onset until the start of Hold 2. Fascicle stretch amplitude was calculated as the peak-to-peak difference in fascicle lengths between the end of Hold 1 and the start of Hold 2. Fascicle velocity was calculated as the maximum derivative of fascicle length over the same period. The mean values of active torque, muscle activity level, and absolute fascicle length during Hold 1 and Hold 2 were quantified for each trial of each condition for both experiments. These three outcome variables, as well as fascicle shortening and stretch amplitudes, the maximum fascicle velocity, the maximum torque matching error (see below), and the coefficient of variation in active torque during Hold 2 were then averaged among all valid trials within each condition and statistically analyzed.

The ability of the participants to match the desired torque trace was assessed by subtracting the active torque from the mean of the two displayed torque traces over a period from contraction onset until the end of Hold 2 (Fig. 1). The ascending ramp was included in this calculation so that fascicle shortening velocities and motor unit discharge rates would likely be similar among conditions (49). The maximum deviation in torque matching was then normalized to the MVC torque. Trials with torque matching errors of less than 10% MVC were included in the analysis and considered as valid. Valid trials were also required to have active torques between the end of Hold 1 and start of Hold 2 that were not more than 0.8 Nm (Experiment 1) or 2 Nm (Experiment 2) below the mean active torque during Hold 2. Values of 0.8 Nm and 2 Nm were chosen because pilot testing revealed that the mean torque variability among conditions during Hold 2 was 0.6±0.1 Nm (Experiment 1) and 0.8±0.6 Nm (Experiment 2), and we opted for a threshold that was two standard deviations above these mean torque variabilities.

### Statistics

Statistical analysis was performed in GraphPad Prism (9.1.2 64-bit version, San Diego, California, USA) with a 5% (two-tailed) alpha level. Two-way repeated-measures mixed-effects analyses with a Greenhouse-Geisser correction were performed to identify mean differences among conditions and hold phases in active torque, muscle activity level (normalized to MVC), and absolute fascicle length. One-way repeated-measures mixed-effects analyses with a Greenhouse-Geisser correction were performed to identify mean differences among conditions in fascicle shortening amplitude, fascicle stretch amplitude, maximum fascicle stretch speed, and the coefficient of variation in active torque during Hold 2. Following a significant effect or interaction, individual-variance-based Holm-Sidak multiple comparisons were performed among conditions. Data are presented as mean ± SD.

## RESULTS

### Experiment 1: Partial muscle deactivation at different rates

#### Data exclusion

Results from Experiment 1 are based on *n*=11 (5 women, age: 26±2 yr; mass: 72±14 kg; height: 176±11 cm). One participant was to unable to match the desired torque reduction rate in the High-Mod conditions and did not complete the experimental session, and one participant had no valid trials in the Fast condition. Fascicle kinematics from two additional participants were excluded because of inaccurate automated fascicle tracking based on visual inspection.

#### Torque matching

A total of 21±1 (range: 19-23) submaximal voluntary contractions per participant were recorded and 16±2 (range: 13-19) were valid. At least 2±1 (Rapid), 3±1 (Fast), and 4±1 (Ref, Slow, Med) valid trials per condition were recorded between participants. The maximum active dorsiflexion torque at 10±2° plantar flexion was 41.0±9.3 Nm (range: 29.9-62.4 Nm). The torque matching (Fig. 2A) in the valid trials was significantly better in Ref (5.2±0.8% MVC, *p*≤.007) than the other conditions (Slow: 6.7±1.0%; Med: 6.4±1.1%; Fast: 6.8±1.0%; Rapid: 7.5±1.1%).

**Figure 2.**
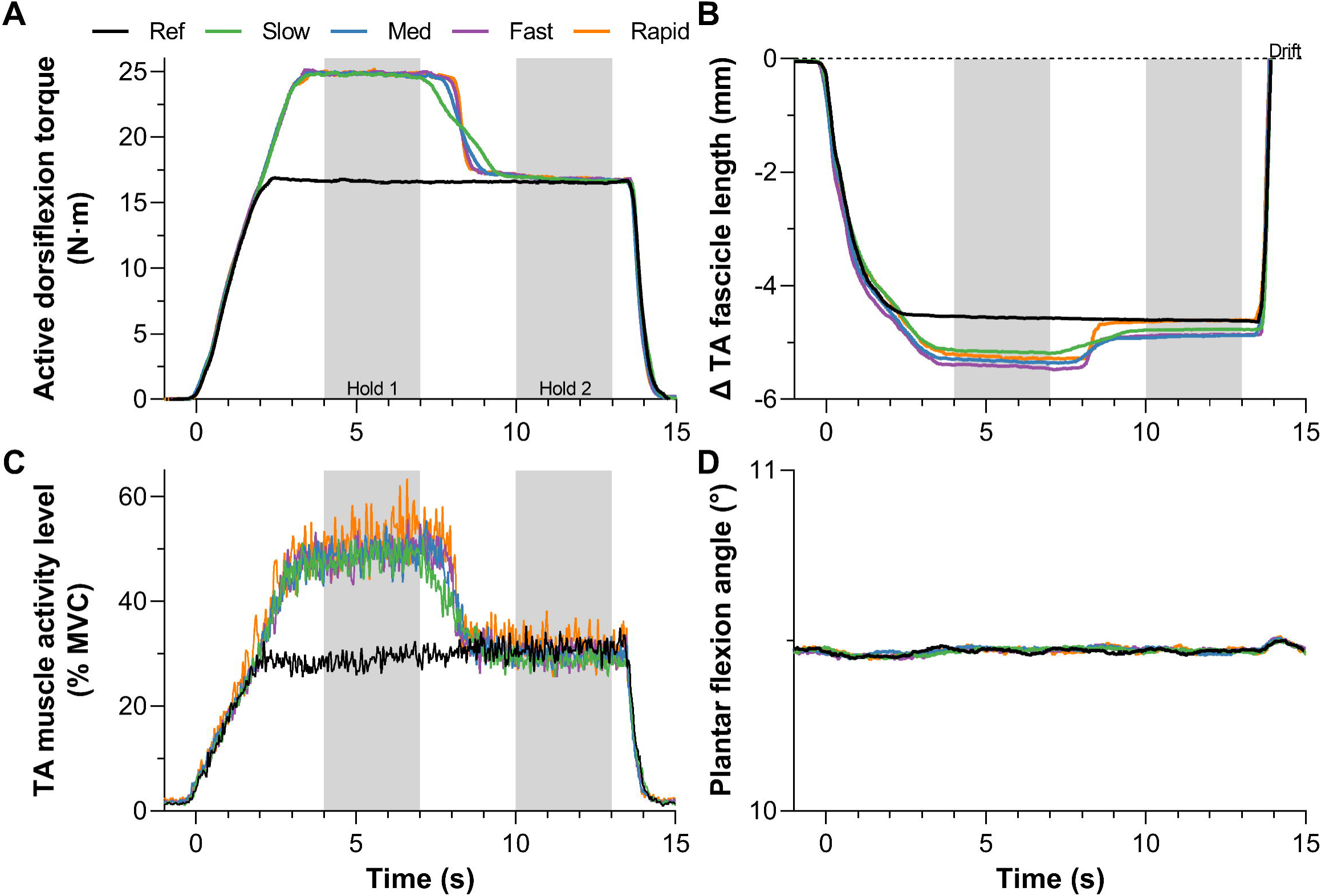
Mean (*n*=11) (A) active torque-time, (B) tibialis anterior (TA) fascicle length change-time (*n*=10 only), (C) TA muscle activity level-time, and (D) crank arm angle-time traces during the five fixed-end contraction conditions. The conditions included one reference (Ref) condition at a moderate torque level and four conditions at a high then matched moderate (High-Mod) torque level. The High-Mod conditions differed in the rate of decreased torque, which was either slow, medium (Med), fast, or rapid. Muscle activity levels are presented as rectified EMG signals that were smoothed with a second-order zero-lag Butterworth filter at 10 Hz and normalized to maximal voluntary contraction. Fascicle length changes are relative to the passive fascicle length prior to contraction onset and optical-flow-based tracking drift caused longer lengths after than before contraction. The grey shaded areas indicate the two analyzed steady-state phases (4-7 s and 10-13 s) of the contractions. Note that the histories of fascicle length change to arrive at the same torque level are different.

#### Hold 1

As expected, the active torques and TA muscle activity levels (Fig. 2A&C) during Hold 1 were significantly higher in the High-Mod conditions compared with Ref (Δ8.1 to 8.3 Nm [Δ19.9 to 20.1% MVC], *p*<.001; Δ19.5 to 22.8% MVC, *p*<.001, respectively), but not significantly different among High-Mod conditions (Δ0.0 to 0.1 Nm, *p*=.940; -0.3 to 3.3% MVC, *p*≥.540; Table 1). The fascicle lengths (Fig. 2B) over the same period were also significantly shorter in the High-Mod conditions compared with Ref (Δ-0.8 to -2.2 mm, *p*≤.049; note that one value was excluded from the Rapid condition), but not significantly different among High-Mod conditions (Δ0.1 to 1.4 mm, *p*≥.176; Table 1).

**Table 1.** Changes in normalized active torque, normalized tibialis anterior (TA) muscle activity level, and TA fascicle length from contraction onset to the first hold phase (Hold 1) and from Hold 1 to the second hold phase (Hold 2) among the five fixed-end conditions in Experiment 1

|  | Contraction onset to Hold 1 |  |  | Hold 1 to Hold 2 |  |  |
| --- | --- | --- | --- | --- | --- | --- |
| | $\Delta$ Torque (% MVC) | $\Delta$ EMG (% MVC) | $\Delta$ Length (mm) | $\Delta$ Torque (% MVC) | $\Delta$ EMG (% MVC) | $\Delta$ Length (mm) |
| Reference | 40.3 (1.4)* | 33.9 (12.1)* | -4.6 (1.3)* | -0.2 (0.3) | 2.0 (1.7) | -0.1 (0.1) |
| Slow | 60.1 (1.0) | 53.5 (16.4) | -5.3 (1.4) | -19.4 (0.6) | -19.6 (7.8) | 0.4 (0.2) |
| Medium | 60.3 (1.1) | 53.8 (16.4) | -5.5 (1.4) | -19.5 (0.7) | -19.1 (6.5) | 0.4 (0.2) |
| Fast | 60.3 (1.6) | 53.5 (16.1) | -5.5 (1.2) | -19.5 (0.8)# | -18.9 (6.3) | 0.5 (0.3) |
| Rapid | 60.4 (2.0) | 56.7 (21.1) | -5.4 (2.0)<sup>§</sup> | -19.5 (0.9) | -20.7 (10.2) | 0.6 (0.3)<sup>§</sup> |

Values are means (standard deviations); $n = ^\S 10$ or 11. \*Significantly different to other conditions ( $P \leq 0.05$ ). #Significantly different to Reference. Comparisons made among conditions by two-tailed two-way repeated-measures mixed-effects models with Holm-Sidak post hoc correction.

#### Descending ramp

The fascicle stretch amplitudes (Table 1 and Fig. 3D) and velocities were significantly higher in the High-Mod conditions compared with Ref (Δ0.4 to 0.7 mm, *p*≤.001; Δ0.9 to 5.2 mm·s^-1^, *p*≤.011), as expected. The fascicle stretch amplitudes were, however, significantly less in Slow compared with both Fast (*p*=.035) and Rapid (*p*=.001), but not compared with Med (*p*=.074). Additionally, the fascicle stretch amplitudes were significantly less in Med relative to Rapid (*p*=.002), but not significantly different between Med and Fast (*p*=.060) or between Fast and Rapid (*p*=.220). The fascicle stretch velocities were significantly different among all High-Mod conditions, which indicates that the different torque reduction rates effectively modulated the rate of fascicle stretch (Δ0.6 to 4.3 mm·s^-1^, *p*≤.041).

**Figure 3.**
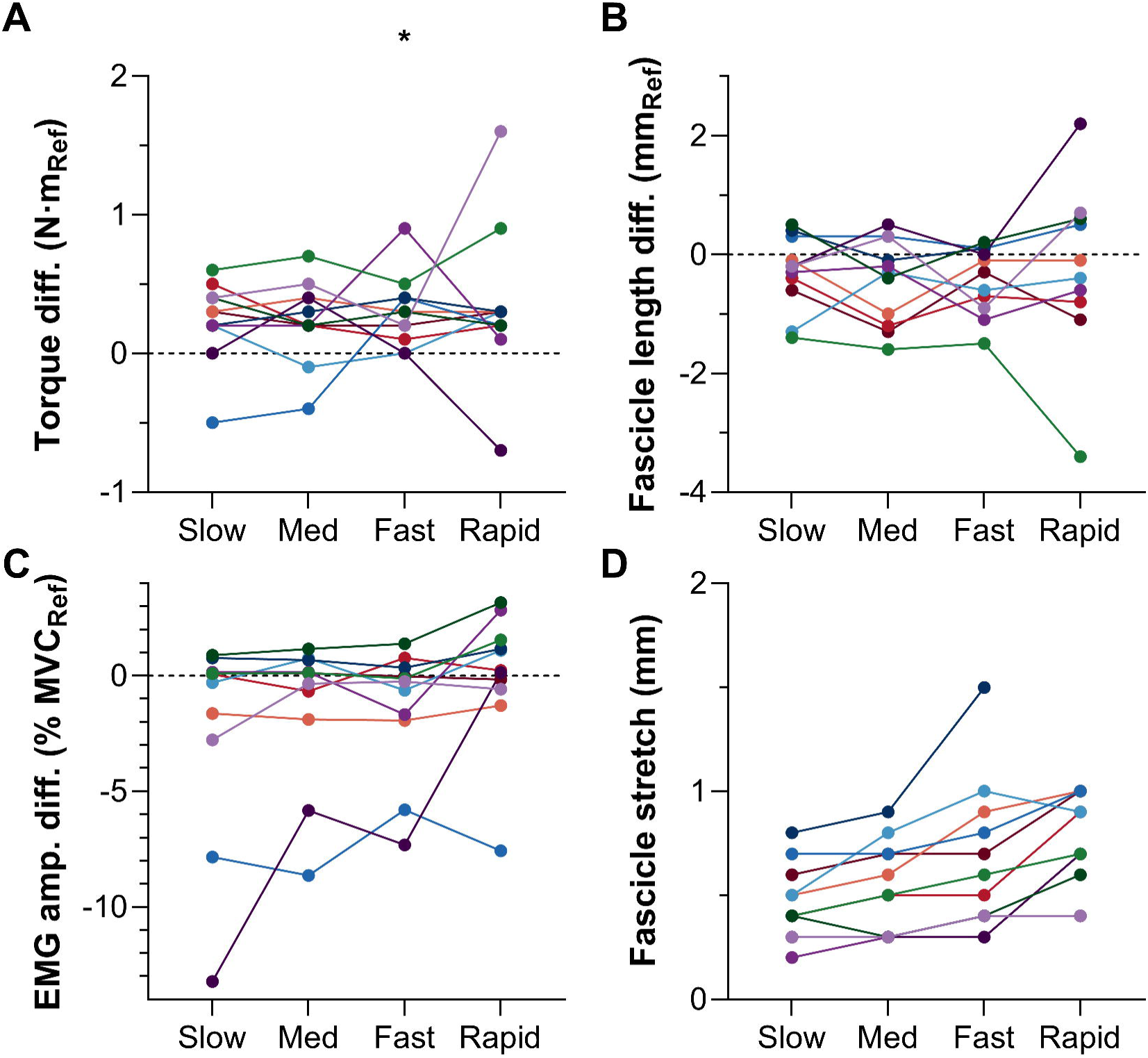
Individual differences for (A) active torque, (B) tibialis anterior (TA) fascicle length, and (C) TA muscle activity level during Hold 2, and for (D) TA fascicle stretch amplitude from Hold 1 to Hold 2 of the four high to moderate (High-Mod) torque level conditions relative to the reference (Ref) condition. The asterisk in A indicates that mean active torque was significantly higher in the Fast condition than the Ref condition. Each color represents a different participant.

#### Hold 2

As this was a torque-controlled experiment, the active torques during Hold 2 were not significantly different in the High-Mod conditions compared with Ref (Δ0.2 to 0.3 Nm [Δ0.7 to 0.8% MVC], *p*≥.194), except for between Fast and Ref (Δ0.3±1.9 Nm, *p*=.031; Table 1 and Fig. 3A). However, this difference falls within the measurement error of the torque sensor. The coefficient of variation in active torque during Hold 2 was not significantly different among conditions including Ref (1.4 to 1.7% [0.6±0.1 Nm], *p*≥0.083). The TA muscle activity levels during Hold 2 were also not significantly different in the High-Mod conditions relative to Ref (Δ-2.2 to 0.6% MVC, *p*≥.607; Table 1 and Fig. 3C). Additionally, the fascicle lengths over the same period were not significantly different in the High-Mod conditions compared with Ref (Δ-1.5 to -0.3 mm, *p*≥.249; Table 1 and Fig. 3B).

### Experiment 2: Partial muscle deactivation of different amounts

Results from Experiment 2 are based on *n*=8 (3 women, age: 25±1 yr; mass: 72±10 kg; height: 178±8 cm). One participant was to unable to match the desired torque reduction traces in any Max condition and did not complete the experiment session. One participant was excluded because the ultrasound images and analog data could not be synchronized. Fascicle kinematics from two additional participants were excluded because of inaccurate automated fascicle tracking based on visual inspection.

#### Torque matching

A total of 21±3 (range: 17-24) submaximal voluntary contractions per participant were recorded and 16±3 (range: 14-21) were valid. At least 2±1 (Max-Min, Max-Low, Max-Mod), 3±1 (Mod), and 4±1 (Min, Low) valid trials per condition were recorded between participants. The maximum active dorsiflexion torque at 10±0° plantar flexion was 41.4±9.6 Nm (range: 28.3-54.9 Nm). The torque matching (Fig. 4A) in the valid trials was significantly better in the reference trial (Min: 4.0±0.7% MVC; Low: 4.4±0.9% MVC; Mod: 5.3±1.2% MVC, *p*<.001) than the respective Max condition (Max-Min/Low/Mod: 8.4±1.0/7.9±0.9%/8.0±1.2% MVC).

**Figure 4.**
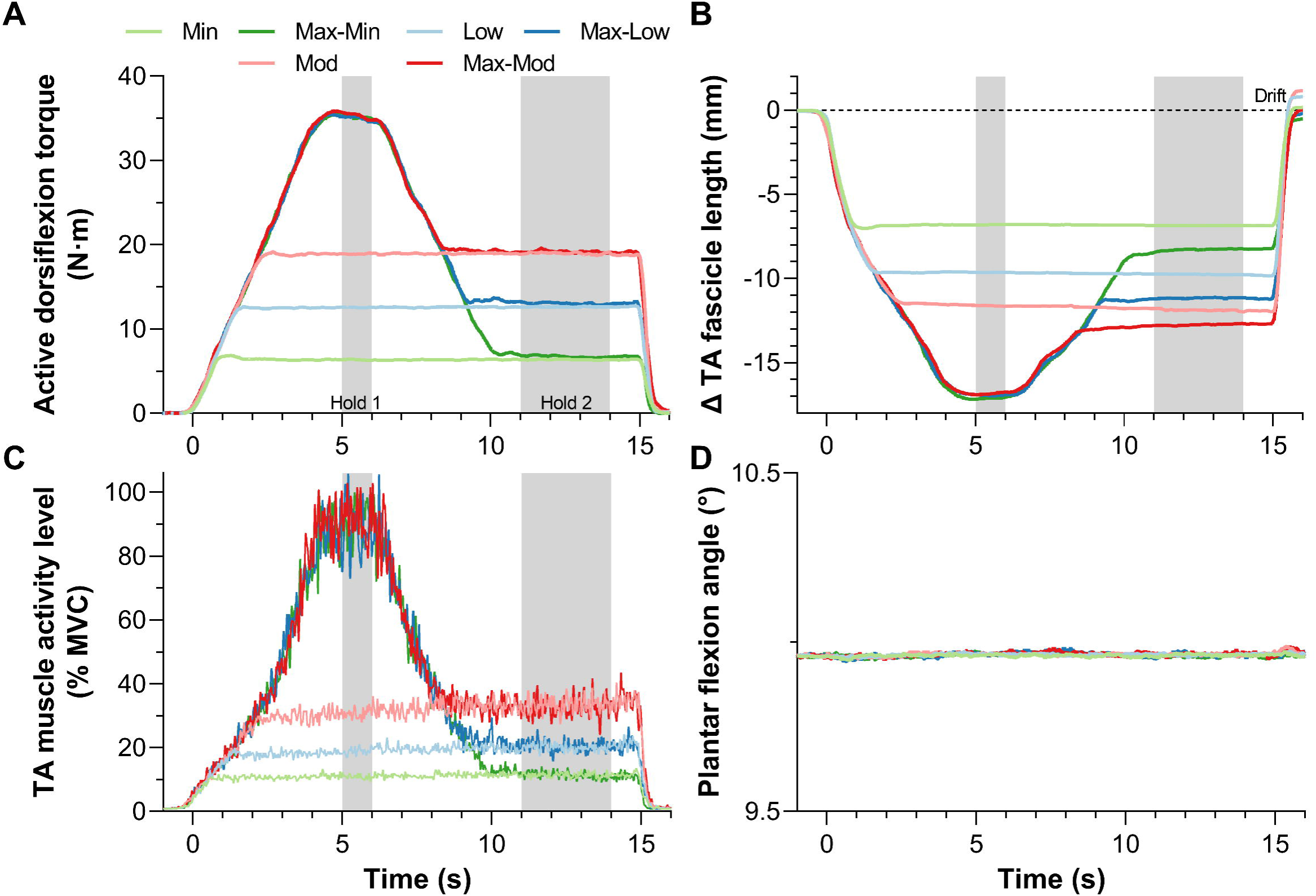
Mean (*n*=8) (A) active torque-time, (B) tibialis anterior (TA) muscle fascicle length change-time (*n*=7 only), (C) TA muscle activity level-time, and (D) crank arm angle-time traces during the six fixed-end contraction conditions. The conditions included three reference conditions at minimal (Min), low, and moderate (Mod) torque levels and three conditions at a maximal then matched minimal (Max-Min), low (Max-Low), or moderate (Max-Mod) torque level. The latter three conditions thus differed in the activation level reduction. Muscle activity levels are presented as rectified EMG signals that were smoothed with a second-order zero-lag Butterworth filter at 10 Hz and normalized to maximal voluntary contraction. Fascicle length changes are relative to the passive fascicle length prior to contraction onset and optical-flow-based tracking drift caused shorter lengths in the Max versus reference conditions. The grey shaded areas indicate the two analyzed steady-state phases (5-6 s and 11-14 s) of the contractions.

#### Hold 1

As expected, the active torques and TA muscle activity levels (Fig. 4A&C) during Hold 1 were significantly higher in the Max conditions compared with the respective reference condition (Δ16.4 to 28.8 Nm [Δ39.6 to 69.6% MVC], *p*<.001; Δ60.8 to 80.1% MVC, *p*<.001; Table 2). Additionally, these same variables were significantly different among reference conditions (Δ6.2 to 12.6 Nm [Δ14.9 to 30.2% MVC], *p*<.001; Δ8.0 to 19.8% MVC, *p*<.001), but not significantly different among Max conditions (Δ-0.2 to 0.3 Nm [Δ-0.3 to 0.5% MVC], *p*≥.445; Δ-2.3 to 2.8% MVC, *p*=.682). The fascicle lengths over the same period were also significantly shorter in the Max conditions compared with the respective reference condition (Δ-5.2 to - 10.0 mm, *p*<.001; note that one value was excluded from the Max-Min condition; Table 2), as well as significantly different among reference conditions (Δ-2.0 to -4.8 mm, *p*≤.002), but not Max conditions (Δ- 0.1 to 0.1 mm, *p*≥.917).

**Table 2.** Changes in normalized active torque, normalized tibialis anterior (TA) muscle activity level, and TA fascicle length from contraction onset to the first hold phase (Hold 1) and from Hold 1 to the second hold phase (Hold 2) among the six fixed-end conditions in Experiment 2

|  | Contraction onset to Hold 1 |  |  | Hold 1 to Hold 2 |  |  |
| --- | --- | --- | --- | --- | --- | --- |
| | $\Delta$ Torque (% MVC) | $\Delta$ EMG (% MVC) | $\Delta$ Length (mm) | $\Delta$ Torque (% MVC) | $\Delta$ EMG (% MVC) | $\Delta$ Length (mm) |
| Min | 15.5 (0.8)* | 10.9 (2.0)* | -7.2 (1.9)* | 0.0 (0.2)* | 0.6 (0.4)* | 0.0 (0.1)* |
| Max-Min | 85.1 (1.8) | 90.9 (7.2) | -17.4 (3.7)<sup>§</sup> | -68.8 (1.9) | -79.5 (6.7) | 8.8 (1.8)<sup>§</sup> |
| Low | 30.4 (0.8)* | 18.9 (3.3)* | -10.0 (2.6)* | 0.2 (0.3)* | 1.1 (1.2)* | -0.1 (0.1)* |
| Max-Low | 84.8 (2.0) | 88.7 (9.2) | -17.1 (3.8) | -53.2 (2.1) | -68.0 (9.6) | 5.8 (1.3) |
| Mod | 45.7 (0.5)* | 30.7 (4.8)* | -12.0 (3.0)* | 0.2 (0.5)* | 2.7 (1.8)* | -0.3 (0.2)* |
| Max-Mod | 85.3 (1.9) | 91.5 (8.8) | -17.0 (3.5) | -38.9 (2.1) | -58.9 (7.7) | 4.1 (1.1) |

Values are means (standard deviations); $n = ^\S 7$ or 8. \*Significantly different to respective Max condition. Comparisons made among conditions by two-tailed two-way repeated-measures mixed-effects models with Holm-Sidak post hoc correction.

#### Descending ramp

The fascicle stretch amplitudes (Table 2 and Fig. 5D) and velocities were significantly higher in the Max conditions compared with the respective reference condition (Δ3.8 to 8.6 mm, *p*<.001; Δ5.8 to 9.1 mm·s^-1^, *p*<.006), as expected. Among Max conditions, the fascicle stretch amplitudes were significantly different (Max-Min/Low/Mod: 8.8±1.9/6.0±1.5/4.1±0.9 mm; *p*≤.001), but the fascicle velocities were not significantly different (Max-Min/Low/Mod: 9.8±4.6/8.6±2.8/7.8±1.7 mm·s^-1^; *p*=.428), which indicates that the different torque reduction amplitudes effectively modulated the amount of fascicle stretch.

**Figure 5.**
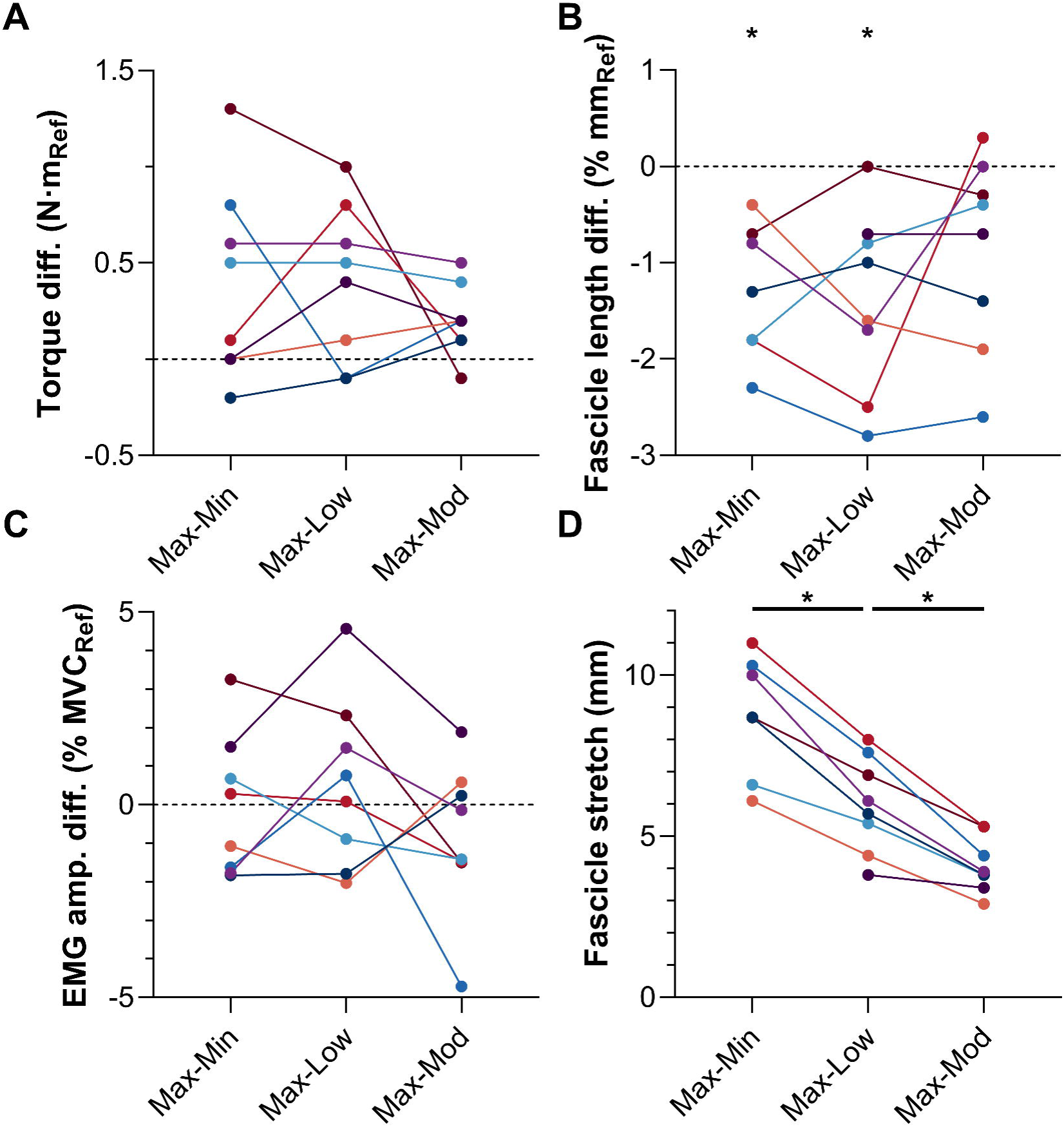
Individual differences for (A) active torque, (B) tibialis anterior (TA) fascicle length, and (C) TA muscle activity level during Hold 2, and for (D) TA fascicle stretch amplitude from Hold 1 to Hold 2 of the three maximal (Max) to lower torque level conditions relative to the respective reference condition (i.e. minimal [Min], low, or moderate [Mod]). The asterisks in B indicate that mean fascicle lengths were significantly shorter in the Max conditions than the respective reference conditions, and the asterisks in D indicate that mean fascicle stretch amplitudes significantly decreased from Max-Min to Max-Low and from Max-Low to Max-Mod. Each color represents a different participant.

#### Hold 2

As this was a torque-controlled experiment, the active torques were not significantly higher in the Max conditions compared with the respective reference condition (Δ0.2 to 0.4 Nm [Δ0.5 to 0.9% MVC], *p*≥.054; Fig. 5A). The coefficient of variation in active torque during Hold 2 (Fig. 6A) was also significantly higher in Max-Min (3.9±0.8% [0.6±0.1 Nm], *p*=.036; Fig. 6B) versus Min (2.7±0.8% [0.4±0.1 Nm]) and Max-Low (2.6±0.8% [0.8±0.2 Nm], *p*=.031) versus Low (1.9±0.4% [0.6±0.1 Nm]). However, the difference was not significant between Max-Mod (2.1±0.4% [1.0±0.2 Nm], *p*=.988) versus Mod (2.1±0.5% [1.0±0.3 Nm]).

**Figure 6.**
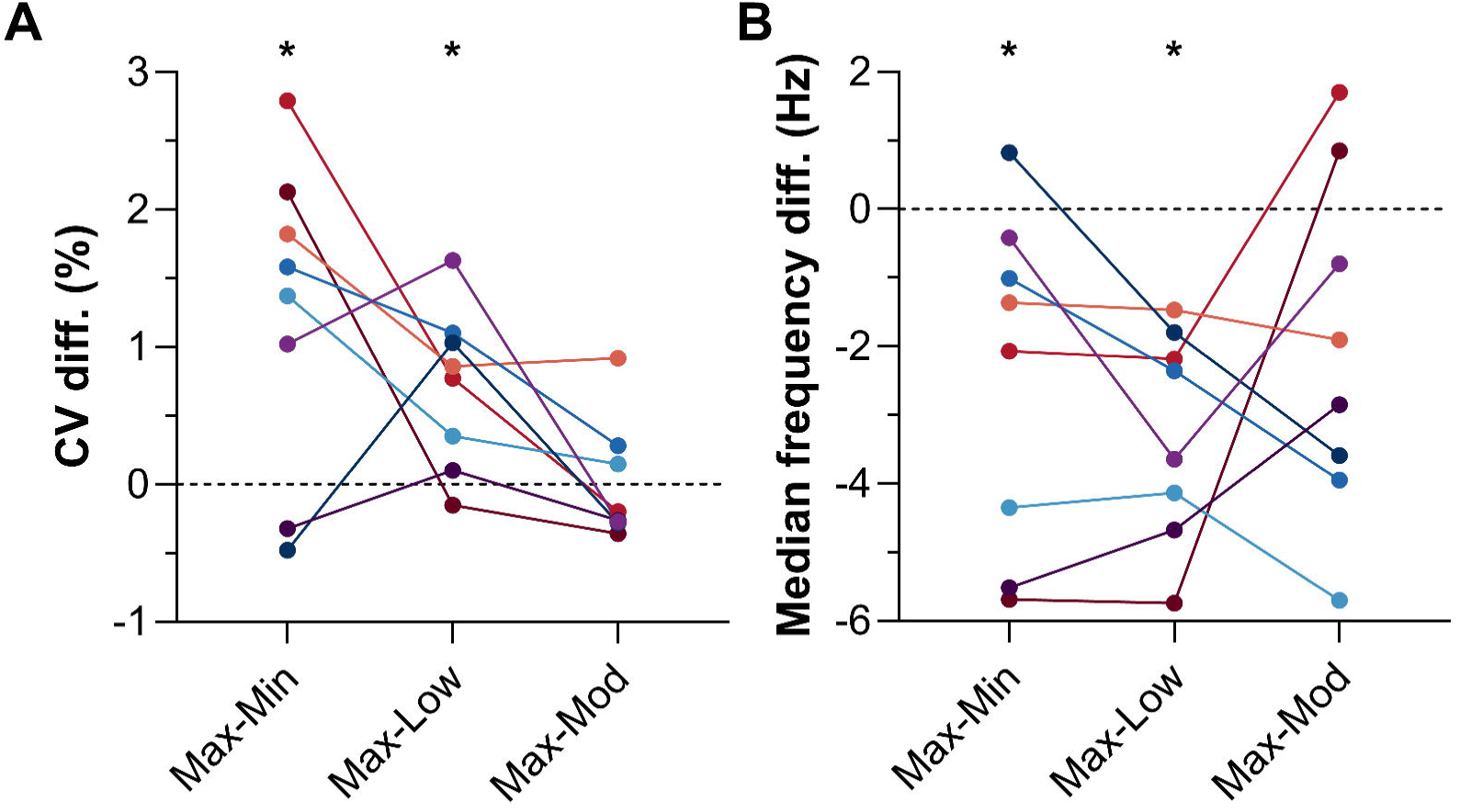
(A) Individual differences (diff.) for the coefficient of variation (CV) in active torque and the median frequency of tibialis anterior’s EMG signal during Hold 2 of the three maximal (Max) to lower torque level conditions relative to the respective reference condition (i.e. minimal [Min], low, or moderate [Mod]). The asterisks in A and B indicate that mean CVs and median frequencies were significantly higher and lower, respectively, in the Max conditions than the respective reference conditions. Each color represents a different participant.

Consequently, we also explored whether there were mean differences between each Max condition and its respective reference condition for the coefficient of variation in muscle activity level and the median frequency of the bandpass filtered EMG signal during Hold 2. The coefficient of variation in muscle activity level was significantly higher in Max-Min versus Min (Δ3.8±2.0%, *p*=.003), but not significantly different between Max-Low versus Low (Δ2.6±3.1%, *p*=.093) and Max-Mod versus Mod (Δ-0.6±2.9%, *p*=.580). The median frequency (Fig. 6B) was significantly lower in Max-Min (54.4±8.2 Hz, *p*=.0496) versus Min (56.9±7.0 Hz) and Max-Low (54.7±7.2 Hz, *p*=.0016) versus Low (58.0±6.9 Hz), but not significantly different between Max-Mod (57.2±6.9 Hz, *p*=.0558) versus Mod (59.2±6.7 Hz).

The TA muscle activity levels during Hold 2 were not significantly different between the comparisons of interest (Δ-0.8 to 0.6% MVC, *p*≥.631; Fig. 5C). However, the fascicle lengths during Hold 2 were significantly different among two of three of the comparisons of interest (Max-Min vs. Min: Δ-1.1±0.7 mm, *p*=.019; Max-Low vs. Low: Δ-1.4±0.9 mm, *p*=.013; Max-Mod vs. Mod: Δ-0.9±1.0 mm, *p*=.084; Fig. 5B), which was not expected, but these differences were similar in magnitude to the amount of fascicle tracking drift (-0.5 to 1.4 mm; Fig. 4B).

## DISCUSSION

We sought to systematically assess whether fascicle stretch due to elastic tissue recoil during partial muscle deactivation can trigger rFE-related mechanisms. rFE was indirectly assessed by comparing TA’s steady-state EMG amplitude (i.e. muscle activity level) following fascicle stretch with a reference condition without fascicle stretch in torque-controlled experiments. It was expected: 1) that rFD would increase with increasing activity level (40) and fascicle shortening amplitude (11, 19, 39) at a constant joint angle, and: 2) that rFE would more than offset the additional rFD from initially higher-force contractions, resulting in a lower muscle activity level following fascicle stretch as previously shown in torque-controlled experiments, which has been termed ‘activation reduction’ (35–37). However, we found that different fascicle stretch rates and fascicle stretch amplitudes did not significantly affect TA’s steady-state muscle activity level with respect to the reference conditions at similar active dorsiflexion torques. This potentially indicates that fascicle stretch from partial muscle deactivation triggered rFE-based mechanisms that precisely offset the additional rFD developed during the higher-force ascending ramp. Intriguingly, rFE-based mechanisms that implicate the engagement of a parallel elastic element (12, 13) might help to explain the increased torque variability we observed during Hold 2 following fascicle stretch because of increased passive force contributions that increase the total force output of each motor unit. Correspondingly, we found some indirect evidence for changes to TA’s voluntary control following larger fascicle stretches (see below). Although our interpretations require independent verification, our results alone imply that the history of force production during fixed-end contractions affects the subsequent control of muscle force more than the steady-state neuromechanical output.

Our primary goal was to test whether stretch-induced rFE can reduce the muscle activity level at a given joint torque. *In vivo*, rFE manifests itself following MTU stretch as either an increased steady-state torque about a joint at a given muscle activity level or a reduced muscle activity level at a given joint torque (i.e. presumed muscle force). Fascicle stretch during MTU stretch was sometimes quantified (26–29) but typically assumed (50–52), which is problematic because fascicle shortening and thus shortening of the muscle’s passive elastic elements can occur with increasing activation during MTU stretch (10, 11, 23), leading to reduced rFD despite appearing as rFE. This study is different to most previous *in vivo* rFE studies because we did not stretch the MTU to induce fascicle stretch, but we induced fascicle stretch alone via elastic tissue recoil due to partial muscle deactivation at a constant MTU length, similar to Oskouei and Herzog (33). In Experiment 1, we found that different rates of partial muscle deactivation resulted in fascicle stretch rates of between 1.6 and 5.9 mm·s^-1^. These differences for fascicle stretch rate were not expected to affect TA’s steady-state muscle activity level following stretch based on findings from cat soleus muscle and the human adductor pollicis, which showed rFE was not significantly different following stretch rates of 2 to 32 mm·s^-1^ (42) and following thumb rotation speeds of 10 to 60°·s^-1^, respectively (34). Similarly, we found that the rate of partial muscle deactivation did not significantly affect TA’s steady-state activity level, but unlike these former studies, we did not find evidence of rFE based on lower steady-state activity levels relative to the refence condition.

The insignificant reduction in muscle activity level following fascicle stretch either indicates that rFE-based mechanisms were not triggered during fascicle stretch or that the rFE following fascicle stretch did not exceed the initial rFD developed during the ascending ramp. As we observed mean fascicle stretch amplitudes of 1 mm and additional mean shortening magnitudes relative to Ref of 1 mm in Experiment 1, we performed Experiment 2. This is because it is unlikely that we had the measurement certainty to detect a presumed increase in rFD of 1-2% during the High-Mod contractions, which is based on our previous findings of 4% less rFD following 2-4 mm reductions in fascicle shortening during MTU stretch (10, 11). Experiment 2 therefore involved larger partial muscle deactivations of varying amplitudes that caused fascicles to stretch more between mean amplitudes of 4 to 9 mm, and the higher initial activity level in Max conditions caused fascicles to shorten 5 to 10 mm more relative to the reference conditions. However, again we did not find significant reductions in muscle activity level following fascicle stretch relative to the respective reference conditions, despite rFD potentially being 5-10% higher in the Max versus reference conditions. It is therefore possible that fascicle stretch during partial muscle deactivation abolished the additional rFD through rFE-based mechanisms.

We are not the first to suggest that rFE-based mechanisms are triggered during a partial drop in muscle force during voluntary fixed-end contractions. Oskouei and Herzog (33) previously implicated rFE-based mechanisms to explain the higher steady-state forces following partial muscle deactivations from 60% or 100% MVC to 30% MVC during EMG-controlled experiments on the human adductor pollicis. However, Oskouei and Herzog (33) suggested that post-activation potentiation rather than stretch of the contractile elements contributed to this rFE, but this is not possible to verify as fascicle kinematics were not quantified in that study. In our experiments, we observed fascicle stretch during partial muscle deactivation, but support for rFE based on lower steady-state muscle activity levels following fascicle stretch was not found. However, we do not believe this finding reflects no rFE. This is because other work on the cat soleus muscle found that rFE was abolished in a dose-dependent manner by the magnitude of preceding shortening (53), which indicates that rFD can mask rFE. Specifically, this work found no clear (0±1%) rFE or rFD when the shortening magnitude exceeded the stretch magnitude by 50% during muscle-tendon unit shortening-stretch cycles (53), which agrees with our findings following larger shortening than stretch amplitudes at the fascicle level of 50 to 90%. Taken together, these findings indicate that shortening-induced rFD abolishes stretch-induced rFE when the shortening amplitude is at least two times the stretch amplitude. Intriguingly though, any rate or magnitude of stretch appears to be sufficient to offset the additional rFD from additional shortening.

Our interpretation that rFE-based mechanisms were triggered during fascicle stretch from elastic tissue recoil during partial muscle deactivation is dependent on our first assumption that increased muscle activity coupled with additional fascicle shortening led to additional rFD. If additional rFD was not induced in the initially higher-force contractions, then rFE-based mechanisms would not need to be triggered during fascicle stretch to cause equivalent neuromechanical output following stretch relative to the reference conditions. This possibility should be kept in mind when interpreting our data considering that rFD was found to be independent on shortening amplitude during submaximal (but not maximal (54)) voluntary contractions (10, 55, 56). Additionally, we tested on the approximate plateau region of TA’s maximal force-length relation (45), where rFE was found to be independent of stretch amplitude and also relatively limited (2-4%) compared with longer lengths on the descending limb (6-14%) in the cat soleus muscle (43) and human knee extensors (26). Consequently, we cannot be certain that there was sufficient rFE to offset any potentially additional rFD in the initially higher-force contractions.

Our interpretation that fascicle stretch during partial muscle deactivation triggered rFE-based mechanisms is also dependent on our second assumption that some fibers were actively stretched. Although we cannot confirm whether active or passive stretch of the imaged fascicles occurred from our ultrasound data, we think it is likely that the fibers that were not de-recruited during partial muscle deactivation were actively stretched. This is because these voluntarily-activated fibers must have been active before and during partial muscle deactivation based on Hennemen’s size principle (57, 58), and because we consistently observed concurrent stretch of all imaged fascicles (i.e. within the superficial and deep compartments) for all participants during the descending ramp. Therefore, it is unlikely that other unimaged fascicles remained isometric during partial muscle deactivation, especially because the location of the ultrasound transducer varied between participants and the chance of not imaging isometric fascicles is very slim. Additionally, we observed proximodistal horizontal movement within TA during both the ascending and descending ramps in Experiment 2, which indicates that TA’s more proximal fascicles consistently experienced length changes before its more distal fascicles. Therefore, it is also unlikely that the earliest-recruited fibers did not continue to shorten once the higher-threshold alpha motoneurons started firing to cause active shortening of their associated fibers.

Despite the high likelihood of active fiber and fascicle stretch, neuromechanical output was not enhanced following stretch relative to the reference conditions, which does not support our a priori hypotheses for either experiment. However, to our surprise, mean fascicle stretch magnitudes from 6 to 9 mm in the Max-Low and Max-Min conditions, respectively, reduced torque steadiness during Hold 2. Additionally, there was a strong exploratory repeated-measures relation between fascicle stretch amplitude during the descending ramp and the subsequent torque steadiness in the Max conditions (*r*rm=.76, 95% CI=.43 to .91, *p*<.001). This could implicate rFE-based mechanisms during fascicle stretch due to elastic tissue recoil considering that we previously found no significant rFE following mean fascicle stretch magnitudes of 2 mm (5) and 4 mm (26) within the human vastus lateralis muscle. rFE is thought to arise from the engagement of parallel viscoelastic elements that increase their passive force contributions during active stretch (9, 24, 25, 59, 60). Increased passive forces could subsequently increase the total force output of each motor unit, making voluntary force control less precise under similar activity levels. The idea that mechanical rather than neural contributions were altered following fascicle stretch from partial muscle deactivation is somewhat supported by our finding of similarly variable muscle activity levels between Max-Low or Max-Mod versus the corresponding reference conditions during Hold 2. However, we also found more variable muscle activity levels in Max-Min relative to Min and a lower median frequency of TA’s EMG signal in Max-Min or Max-Low versus the corresponding refence conditions. This latter finding suggests that motor units probably discharged at lower rates during Hold 2, which could have also reduced torque steadiness. However, this possibility requires independent verification from intramuscular and/or high-density EMG recordings under similar contraction conditions.

Rather than or in addition to rFE-based mechanisms reducing torque steadiness, a larger partial deactivation might make it more difficult to subsequently control torque because of plateau potential activation from persistent inward currents (PICs) (61–65). PICs depend on the level of monoaminergic input (particularly serotonin and norepinephrine) and can amplify and prolong excitatory synaptic input to motoneurons, triggering long-lasting self-sustained firing (61). As we found decreased torque steadiness following partial muscle deactivations of 55% MVC and above, this might indicate that plateau potentials from PICs contributed more to force production and reduced torque steadiness at lower muscle activity levels. That is, the initially higher synaptic input during the initially higher-force contractions could have activated plateau potentials that kept some higher-threshold motor units recruited during and following the partial muscle deactivation. The increased motor unit recruitment should have subsequently resulted in a higher muscle activity level relative to the reference contractions, but as this was not observed here, it is possible that the overall discharge rate of the active motor unit pool was also lower following the partial muscle deactivation, which is supported by our median frequency results. Therefore, increased motor unit recruitment from PICs coupled with lower overall discharge rates could additionally or alternatively help to explain the reduced torque steadiness following the partial muscle deactivation. Although this possibility is speculative (66) and cannot be explored with our data, previous findings showing lower discharge rates coupled with less motor unit recruitment following MTU stretch provide partial support only, which indicates that further research on the neural underpinnings of muscle force control is required.

### Limitations

Our results should be interpreted with the following limitations in mind. This study was only powered to detect large effects (*d*z of 1) with 80% power and therefore does not provide conclusive evidence that the history of force production does not affect the subsequent steady-state neuromuscular output during fixed-end contractions. We also did not estimate rFD, but assumed that rFD was present during fixed-end contractions based on previous experiments on the same muscle group within the same laboratory under similar conditions (10, 11). Further, we assumed that rFD would increase with increasing muscle activity coupled with additional fascicle shortening based on previous animal findings (19, 39) and increased rFD of 4-9% during higher-force (25%) shortening contractions of the electrically stimulated human adductor pollicis (40). We also assumed that active stretch of some fibers occurred during partial muscle deactivation despite the inability to confirm this with our ultrasound data. It is possible that antagonistic muscles, such as the triceps surae, contributed to the recorded net joint torque and TA’s EMG signal through co-contraction and cross-talk, but we think that systematic differences between conditions were unlikely and negligible based on previous work (67, 68). Similarly, changes in synergistic activation between conditions were probably unlikely because we observed minimal fatigue during and after the contractions (Table 1 and 2) and TA fascicle lengths were similar at similar torques and activity levels, except in Experiment 2. However, the differences for fascicle length in this experiment were more likely to be due to fascicle tracking drift than to activation differences, which is a limitation of our utilized optical-flow-based tracking method.

### Perspectives and Significance

Our findings indicate that initially high-force contractions become harder to control following larger fascicle stretches and partial muscle deactivations, which implicates rFE-based mechanisms and/or PICs as contributors to reduced muscle force control. Implementing a restricted experimental protocol (e.g. Max-Min and Min conditions only) at relatively longer muscle lengths to increase the chance of inducing rFE (26, 43), in combination with ultrasound imaging and transparent-grid high-density EMG recordings (69), might therefore be worthwhile to better understand the neuromechanical contributors to impaired muscle force control. Similar experiments in people with suspected neuromuscular disorders could be useful to further probe if neuromechanical parameters are useful markers for predicting impairment or monitoring disease progression.

### Conclusions

Contrary to our expectations, we found that fascicle stretch due to elastic tissue recoil during partial muscle deactivation did not subsequently enhance the TA’s steady-state neuromechanical output as reflected by its similar steady-state muscle activity level relative to torque-controlled fixed-end reference contractions. Although this finding in isolation might indicate that rFE-based mechanisms were not triggered by fascicle stretch, this interpretation neglects earlier work showing that rFE was masked by rFD when the preceding shortening exceeded stretch (53). Consequently, it is possible that shortening-induced rFD masked stretch-induced rFE during our experiments and that stretch-induced rFE contributed to the reduced torque steadiness we found following larger fascicle stretch magnitudes and partial muscle deactivations. As the reduced torque steadiness was not fully explained by differences in the amount or variability in TA’s EMG signal, but also by differences in its frequency content, the neural (e.g. plateau potential activation from PICs) and mechanical mechanisms underpinning impaired muscle force control following larger torque drops during fixed-end contractions are worthy of further investigation.

## DATA AVAILABILITY

The raw data from this study will be made available upon publication.

## SUPPLEMENTAL MATERIAL

Links to the raw data will be shared upon publication.

## ACKNOWLEDGMENTS

We thank Leon Lauret (Bochum, North Rhine-Westphalia) for help with the fascicle tracking.

## AUTHOR CONTRIBUTIONS

BJR and DH conceived and designed research, RDL and MK performed experiments, BJR analyzed data, all authors interpreted results of experiments, BJR prepared figures, BJR drafted manuscript, all authors edited and revised manuscript, all authors approved final version of manuscript.

